# Efficient peripheral nerve firing characterisation through massive feature extraction

**DOI:** 10.1101/508341

**Authors:** Carl H Lubba, Ben D Fulcher, Simon R Schultz, Nick S Jones

## Abstract

Peripheral nerve decoding algorithms form an important component of closed-loop bioelectronic medicines devices. For any decoding method, meaningful properties need to be extracted from the peripheral nerve signal as the first step. Simple measures such as signal amplitude and features of the Fourier power spectrum are most typically used, leaving open whether important information is encoded in more subtle properties of the dynamics. We here propose a feature-based analysis method that identifies changes in firing characteristics across recording sections by unsupervised dimensionality reduction in a high-dimensional feature-space and selects single efficiently implementable estimators for each characteristic to be used as the basis for a better decoding in future bioelectronic medicines devices.

## I. INTRODUCTION

Bioelectronic medicines [1] modulate the activity patterns on peripheral nerves by implanted devices. They form a new way of treatment with promise for many conditions such as hypertension and tachycardia [2], [3], sleep apnea [4], rheumatoid arthritis [5] and many more. Today’s devices are still very simple, however, and mostly operate in an open-loop fashion that is not aware of the current activity on the nerve. For future bioelectronic medicines, closed-loop systems that diagnose the signals on target nerves and only block or stimulate when necessary could be much more efficient and effective. It is thus vital for the progress of the field to investigate ways of continuously or periodically characterising peripheral nerve activity and associating its patterns with the physiological state (‘decoding’) to modulate adaptively.

A first step of such a decoding will be the extraction of meaningful properties from the peripheral nerve recording to then associate with the physiological parameters we seek to estimate. Peripheral nerve recordings possess a low signal-to-noise ratio, due to the weak (~10mV) potentials caused by the axons and the presence of other sources of electricity such as muscles. Spatial recording resolution of current non-invasive interfaces (cuff electrodes) is limited as well, making it impossible to differentiate single fibers. The recording is thus made up by compound action potentials (CAP): the superposed activity of many axons in a nerve-bundle. Given these constraints on data quality, decoding is most often based on the amplitude of the rectified and integrated signal [6], [7] or the power (as square of amplitudes from a Fourier spectrum) in a certain frequency band [8].

But are those simple summary values (amplitude, power) sufficient to capture the entire information contained in peripheral nerve recordings or are better measurements possible? Many subtleties are known to exist in peripheral firing such as active fibre diameters, active fascicles and different rhythms [9], many of which will be informative for decoding. If we could identify some of those firing characteristics in a peripheral nerve recording and estimate them in each new observation, we might have a better starting point for decoding. So how can we decide which are the main varying firing characteristics and how can we estimate them – keeping in mind that this estimation has to be very energy-efficient to not deplete the small battery of the implanted device?

One possibility of characterising time series such as peripheral nerve recordings by their dynamical properties is a set of global time series features, of which many have been developed in different disciplines over the past decades [10]. Representing a time series by its dynamical properties using features proved useful for e.g., classification [11], clustering, and forecasting [12]. In this work we seek to leverage the vast literature on time-series features to find better representations of peripheral nerve firing.

Building on a diverse set of over 7 500 time series features [13], [14], we propose an unsupervised analysis method that fulfills two purposes. It (1) automatically infers the types of properties across which peripheral nerve recordings vary most and (2) proposes estimators that quantify these properties (‘characteristics’) in new data. We demonstrate the utility of our approach on simulated data in which the activity characteristics firing rate, myelination ratio, and burstiness were uncovered successfully. The selected estimators for the main peripheral firing characteristics can be efficiently implemented for on-line summarisation of peripheral nerve recordings and thus as a basis for a more accurate decoding in closed-loop bioelectronic medicines.

## II. METHOD

### A. Time-series features

We want to analyse peripheral nerve recordings by their dynamical properties based on a diverse set of estimators for time-series characteristics. To this end we use the Highly Comparative Time Series Analysis (*hctsa*) toolbox [13], [14] which lets us compute more than 7 500 global time-series features measuring e.g., basic statistics of value-distributions, linear correlations, stationarity, entropy and many more. Each feature *f_i_*(*X*) summarises a time series *X* = {*X*_1_, *X*_2_,…, *X_N_*} of length *N* as a single value, *f_i_*: ℝ^*N*^ → ℝ. Using the ensemble of all *M* ≈ 7 500 features *F* = {*f_i_* : *i* = 1,…, *M*}, the *hctsa*-toolbox can transfer our peripheral nerve recordings to data points in an *M*-dimensional feature-space *F*(*X*): ℝ^*N*^ → ℝ^*M*^ for characteristics-based analysis. On the datasets in this study, up to 1500 of our inital 7500 features had special valued outputs and were removed, on average ~6800 remained. Most features of the library are robust against unusual time-series properties, only very short time series with less than ~100 samples will cause a higher percentage to fail. Computation of all features for one of our time series of length 8000 samples took about 340±180s on a single core in a state-of-the-art cluster. If computation time is a constraint, the feature set can be pruned at modest performance decline, see Results III-A.

### B. Detecting the varying characteristics in feature-space

We analyse datasets in the high dimensional feature-space generated by the *hctsa*-toolbox to detect the dynamical properties (‘characteristics’) that varied over time. See Fig. 1 for a schematic overview of our approach. Using *hctsa*, each single time series can be summarised as a point in feature-space; all recordings contained within a dataset form a point-cloud. Features are normalised per dataset by a robust sigmoid transform [13]. If characteristics that vary between recordings are estimated by some of the computed features, their variation will drive the spread of this point cloud along those features. We can thus detect varying signal properties as the main directions of variance using dimensionality reduction methods such as principal component analysis (PCA) or others [15]. These main dimensions make up the low-dimensional space of ‘main varying characteristics’ to project our data into.

**Fig. 1.**
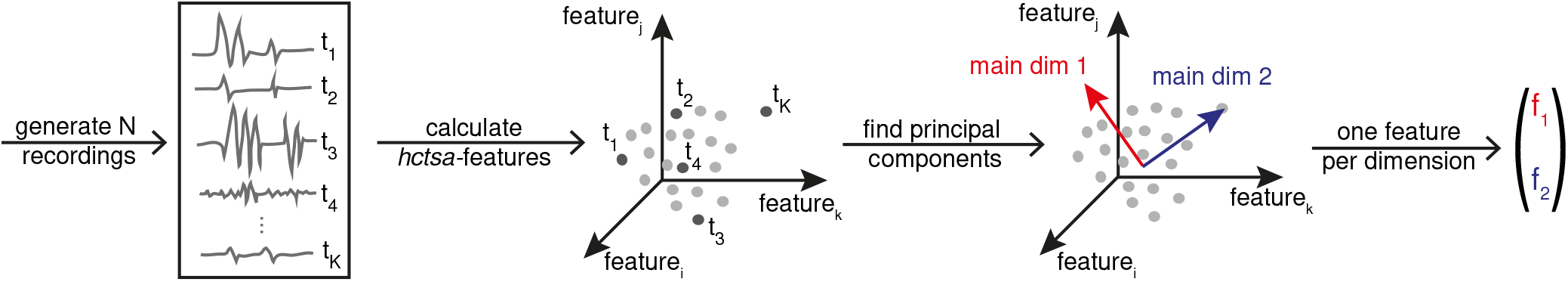
Unsupervised selection of efficient estimators for the main varying signal characteristics. Each of the N peripheral nerve recordings in a given dataset is transferred to a high-dimensional feature-vector using the *hctsa*-toolbox (here visualised as three-dimensional albeit having thousands of dimensions). In property space, we find the main directions of spread in the data caused by the dynamical properties that vary across single recordings. For each component, a single feature can be chosen that estimates one main characteristic.

### C. Selecting efficiently implementable features

So far we have uncovered the main varying signal characteristics across our recording segments as the principal components (PC) in feature-space. Each of these components will depend on the computation of thousands of single features. To efficiently project new data onto the PCs, we approximate each of them by a single representative feature selected by maximum Pearson correlation to the component. In this way we devise our set of efficient estimators for the varying signal characteristics in the dataset at hand.

### D. Datasets

To be in possession of ground truth and to demonstrate the success of our method, we generated surrogate data in the peripheral nerve simulator PyPNS [16]. Four simulated datasets were obtained from a nerve with a fixed length of 5cm containing 500 active axons. For each of the four datasets, we generated 400 to 1000 snippets of length 400ms, sampled at 20kHz, across which two to three of the firing characteristics myelination ratio (0 – 2%), firing rate (0.1 – 10 spikes/ axon/ second), and burstiness (0 – 100% of firing probability imbalance between two alternating intervals) varied uniformly. Our method was then trained to detect these characteristics in the unsupervised manner described above.

## III. RESULTS

### A. Recovery of characteristics

We want to uncover, in a purely data-driven way, the three characterstics firing rate, myelination, and burstiness that varied in our simulator. To this end we transferred all simulated peripheral nerve recordings per dataset into the hctsa-feature-space and found the principal components. Ideally, each of the two to three characteristics varied in a given dataset would be recovered in one principal component (PC) in feature-space each. To demonstrate this correspondence between PCs and characteristics in a first example, Fig. 3 plots the time-series points for the varying characteristics firing rate and burstiness projected onto the first and second PC. As can be seen, firing rate is cleanly captured by the first component, burstiness by the second. At lower firing rates (to the right of the plots in Fig. 3), PC2 is less informative of burstiness, consistent with visual intuition from Fig. 2A and C (burstiness is harder to distinguish for low firing-rate).

**Fig. 2.**
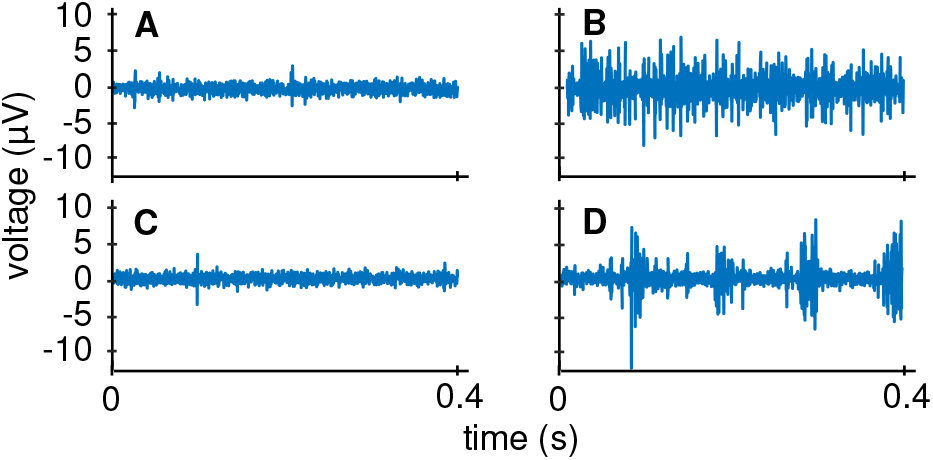
Example simulated recording with varying characteristics firing rate and burstiness. Data was generated in the peripheral nerve simulator PyPNS [16]. (a) Low rate, low burstiness, (b) high rate, low burstiness, (c) low rate, high burstiness, (d) high rate, high burstiness. At low rates, burstiness is not visible even for a human observer in (a) and (c).

**Fig. 3.**
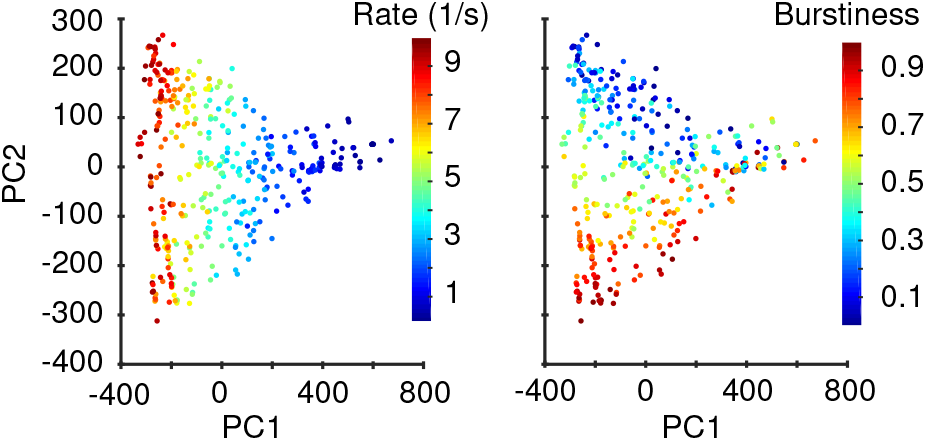
Example for a successful unsupervised detection of the characteristics burstiness and firing rate in the first two principal components (PC) in a normalised *M*-dimensional feature-space. When projecting the time series into the space of the first two principal components as obtained from PCA in our ~6 800-dimensional feature-space (here as axes), each of the two main signal characteristics is captured in one dimension (coloring). Each dot is a time series. At low firing rates (high PC1, to the right of the plot), burstiness cannot be detected anymore as was to be expected from Fig. 2.

As an overview over all our four datasets, Fig. 4A shows the correlation between firing characteristics and each of the first components for datasets with noise-level -6dB (see Fig. 2 for an example recording). On datasets with two varying characteristics, especially in the pairs (bustiness, myelination) and (firing rate, burstiness), each characteristic was perfectly recovered by a single PC each. For the pair (myelination, firing rate), firing rate was not cleanly separated from myelination as noise partly shadowed action potentials from unmyelinated fibres. At three varying characteristics, burstiness and myelination were reasonably captured by the first and second PC but firing rate could not be cleanly separated anymore. Across all datasets, PC1 captured between 60 and 90% of the variance, the PC2 5 to 25%, and PC3 1 to 5%. When randomly selecting subsets of the *hctsa* feature pool to reduce computation time, a characteristics-dependent decline in recovery with modest losses from ~1000 features can be observed in Fig. 4C.

**Fig. 4.**
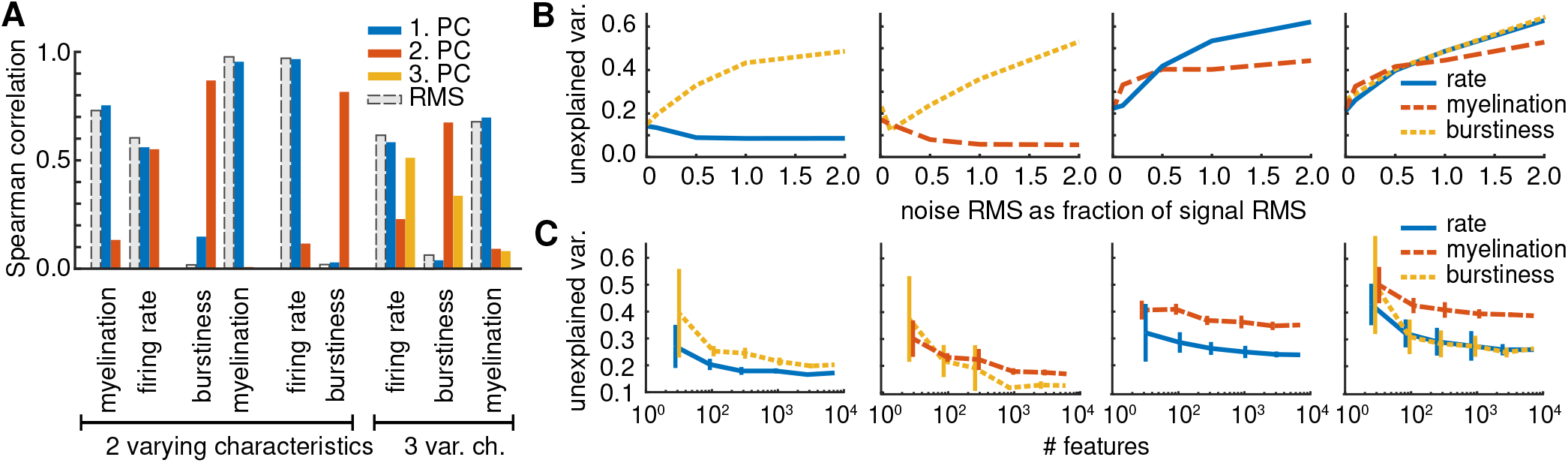
Two varying characteristics are well recovered by PCA in feature space up to a noise-power of half the signal power (−6dB). The second and third principal component provide additional information compared to standard power-measures. (a) Noise RMS was set to half the signal RMS, see Fig. 2 for example time series. Bars indicate the Spearman rank correlation between a single characteristic of a dataset and the classic measure RMS as well as the first principal components (PCs) obtained by dimensionality reduction. RMS behaves largely identical to the first PC. (b) Abscissa is the unexplained variance between the actual signal characteristics and the ones linearly regressed against the first dimensions, Eq. (1). (c) When only considering a subset of all ~6 800 *hctsa*-features, characteristics recovery declines. Each number of features randomly sampled 10 times, SNR=0.1.

### B. Robustness against noise

As a measure of how well the main dimensions obtained by dimensionality reduction captured signal characteristics, we linearly regressed each characteristics *c* as set in the simulation against the first main dimensions **D** (2 main dimensions for 2 varying characteristics, 3 for 3):

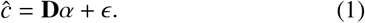

The unexplained variance 1 − *R*^2^(*ĉ, c*) gave an estimation of how well the input characteristics could be retrieved by the output of our method. Until a noise RMS of half the signal RMS (−6dB), the characteristics set in our simulation were recovered well, see Fig. 4B. Myelination ratio was the most robust against noise as expected from the high action potential amplitudes from myelinated fibres, burstiness and firing rate could not be well detected at high noise levels.

### C. What types of features are selected?

So far we have shown that unsupervised dimensionality reduction in feature-space is a promising way of extracting low-dimensional directions of variation in the underlying system that works well in the majority of our simulated neural firing datasets. But can we interpret each dimension in terms of the signal properties that are being measured, and are they sensible given what we know is varying in the underlying system? To answer these questions, we rank single features by their Pearson correlation to the principal components – and therefore the varying characteristics in the data.

In general, the highest Pearson correlations between single features and principal components reached at least 0.8 and often 0.98 or better, meaning that we can find appropriate single features to represent each principle component. For the characteristic myelination ratio, features selected by our method typically evaluate extreme events, outliers, statistics on residuals in local fits. This makes sense as myelinated axons produce very strong peaks. Features corresponding to firing rate components often compute value distribution properties and autocorrelation measures that detect uncorrelated noise vs. natural signals. Burstiness-features measure stationarity and predictability. We therefore automatically identify sensible features.

The selected estimators will be different for every dataset and the purpose of this study is not to select a fixed set of features for real world data from our simulated demonstration datasets. The method has to be rerun for a dataset to analyse as each recording will be different in the composition of fibres, the firing patterns and the interface.

### D. Comparison to standard methods

Our method is able to detect the main varying characteristics in a peripheral nerve recording and select single estimators for them. The unsupervised nature of this characteristics-discovery thus goes beyond unsupervied feature-selection approaches such as Laplacian score [17] or SVD-entropy [18], and further makes use of the most comprehensive feature set available to date.

What do our low dimensional representations add to the standard power-based features computed on peripheral nerve activity? For a brief comparison of our method to RMS as a feature, we added its Spearman rank correlations to the characteristics set in our simulation in Fig. 4A. Naturally, with a univariate power measure, no distinction between different firing characteristics is possible. Interestingly however, our first principal component behaves largely identical with

RMS in terms of correlation to the data characteristics. In the following PCs, more subtle dynamical properties such as burstiness and myelination were recovered. The estimators selected by our method thus cover all signal properties captured by state-of-the-art measures but importantly provide additional information about more subtle firing characteristics.

## IV. CONCLUSIONS

Peripheral nerve decoding algorithms will play an important role in the development of closed loop bioelectronic medicines devices. To date the analysed recordings have been characterised by simple amplitude- or power-based measures. We here demonstrate the feasibility of (1) automatically detecting important signal characteristics and (2) providing simple feature-based estimators for each. Our method is successful on simulated peripheral nerve recordings in which two independent peripheral firing characteristics could be recovered cleanly in most cases. The method provides a low dimensional representation of the data in meaningful dynamical properties that is more informative than the state-of-the-art characterisation by simple power measures. The selected single estimators for important dynamical properties of peripheral firing can be implemented efficiently for the use in next generation bioelectronic medicines devices and the method may find application in related BMI applications as well.

## ACKNOWLEDGMENT

We acknowledge the support by EPSRC grant EP/L016737/1 and EP/N014529/1 as well as Galvani Bioelectronics.

